# High survival following bleaching highlights the resilience of a highly disturbed region of the Great Barrier Reef

**DOI:** 10.1101/2021.10.18.464880

**Authors:** Cathie A Page, Christine Giuliano, Line K Bay, Carly J Randall

## Abstract

Natural bleaching events provide an opportunity to examine how local scale environmental variation influences bleaching severity and recovery. During the 2020 marine heatwave, we documented widespread and severe coral bleaching (75 – 98% of coral cover) throughout the Keppel Islands in the Southern inshore Great Barrier Reef. *Acropora, Pocillopora* and *Porites* were the most severely affected genera, while *Montipora* was comparatively less susceptible. Site-specific heat-exposure metrics were not correlated with *Acropora* bleaching severity, but recovery was faster at sites that experienced lower heat exposure. Despite severe bleaching and exposure to accumulated heat that often results in coral mortality (degree heating weeks ∼ 4 – 8), cover remained stable. Approximately 94% of fate-tracked *Acropora millepora* colonies survived, perhaps owing to reduced irradiance stress from high turbidity, heterotrophic feeding, and large tidal flows that can increase mass transfer. Severe bleaching followed by rapid recovery, and the continuing dominance of *Acropora* populations in the Keppel Islands is indicative of high resilience. These coral communities have survived an 0.8 °C increase in average temperatures over the last 150 years. However, recovery following the 2020 bleaching was driven by the easing of thermal stress, which may challenge their recovery potential under further warming.

**Open Research Statement:** Data are not yet provided but are being compiled. Upon acceptance data will be archived on GitHub.

## Introduction

Global warming has accelerated since 1981 with each new year increasingly likely to rank in the top ten hottest years on record (NOAA, 2021). Oceans are warming, and coral reefs are increasingly experiencing both warmer summers and winters (NOAA, 2021). Marine heat waves bring periods of extreme temperatures and, particularly when combined with high irradiance, can disrupt the functioning of the coral holobiont and lead to bleaching (Baird et al., 2009; Bourne et al., 2008; Brown, 1987; Downs et al., 2013; Hoegh-Guldberg, 1999). The frequency and intensity of marine heat waves, and subsequent coral-bleaching events, are increasing worldwide (Cai et al., 2014; Hoegh-Guldberg, 1999; Hughes et al., 2017a; Timmermann et al., 1999) and have contributed to regional declines in coral cover and diversity (Baker et al., 2008; Bruno and Selig, 2007; De’ath et al., 2012; Gardner et al., 2003; Ortiz et al., 2018; Osborne et al., 2017; van Woesik et al., 2018). As a result, climate change-driven ocean warming is now considered to be the most significant threat to the persistence of globally functioning coral-reef ecosystems (Bellwood et al., 2004; Riegl et al., 2009), and the capacity for corals communities to continue to recover from more frequent and intense coral bleaching is diminishing (Baker et al., 2008; Donovan et al., 2021; Ortiz et al., 2018; van Woesik et al., 2018).

Although bleaching events impact reefs across broad areas, local-scale variability in environmental conditions can lead to heterogeneity in the severity of bleaching and in post-bleaching survival and recovery (Brown, 1987; Fisher et al., 2019; Hoogenboom et al., 2017; Nakamura et al., 2005). Even on the most severely bleached reefs, some coral colonies resist, or recover from bleaching (Hoogenboom et al., 2017; Hughes et al., 2017a, 2019b). Local-scale variation in environmental conditions including tidal movements and wave action (DeCarlo and Harrison, 2019; Donovan et al., 2021; Green et al., 2019), water flow (Nakamura et al., 2005; Nakamura and van Woesik, 2001), turbidity (Fisher et al., 2019; Morgan et al., 2017; Sully and Woesik, 2020), island-shading effects (Fabricius et al., 2004) and cloud cover (Mumby et al., 2001; Skirving et al., 2017) have proven important in driving heterogeneity in bleaching severity and post-bleaching survival within reefs. Quantifying the role that these environmental factors play in driving survivorship and recovery post-bleaching can improve our capacity to predict future coral population trajectories (Donovan et al. 2021).

Variable bleaching responses can also be due to genetic variation with coral populations (Baird et al., 2009; Howells et al., 2013) and their obligate Symbiodiniaceae (*sensu* LaJeunesse et al., 2018) photosymbiont communities (Baker, 2003; Bay et al., 2016). Host sensitivity to heat stress, plasticity in heterotrophy, symbiont community composition and symbiont shuffling can all contribute to within-population variation in bleaching (Abrego et al., 2008; Baker et al., 2008; Bay et al., 2016; Dixon et al., 2015; Grottoli and Palardy, 2006; Howells et al., 2013; Jones et al., 2008; Marangoni et al., 2019). How much of the variability in bleaching during any given heatwave is due to fine scale environmental variation versus within-population genetic variation in corals or their photosymbionts is difficult to quantify. Differential survival of coral genotypes following bleaching and the increased occurrence of heat tolerant *Symbiodinium* clades in surviving corals has also been shown to lead to the directional selection of more heat-tolerant and resilient locally adapted corals (Dixon et al., 2015; Howells et al., 2013; Sully et al., 2019), changing this balance over time.

Coral species also vary in their sensitivity to environmental conditions, including heat and light stress, and are therefore differentially susceptible to bleaching (Guest et al., 2012; Hoogenboom et al., 2017; Loya et al., 2001; Marshall and Baird, 2000; Swain et al., 2017). Many fast-growing branching genera (i.e. *Acropora, Pocillopora, Stylophora* and *Seriatopora*, are often more likely to bleach and die, than slower growing coral taxa (Loya et al., 2001; Marshall and Baird, 2000; Swain et al., 2017; van Woesik et al., 2011). These same branching coral taxa, particularly of the genus *Acropora*, are also vulnerable to other disturbances due to their propensity to fragment following storms and cyclones (Fabricius et al., 2008; Madin, 2005), high disease susceptibility (Patterson et al., 2002; Randall and van Woesik, 2015; Sutherland et al., 2004; Willis et al., 2004), and because they are the preferred prey of corallivores including *Drupella* and crown-of-thorns starfish (Forde, 1992; Pratchett, 2007). In a warmer world in which disease outbreaks, severe tropical cyclones, and regional scale bleaching events are increasing in severity and frequency (Harrison et al., 2019; Harvell et al., 1999; Heron et al., 2016; Hughes et al., 2017a; Ward and Lafferty, 2004; Webster et al., 2005), the ongoing persistence of these sensitive coral taxa is at risk (Côté and Darling, 2010). Certainly the susceptibility of Caribbean acroporids to disease, hurricanes and bleaching has contributed to their endangered listing (Diaz-Soltera, 1999; Precht et al., 2002; Randall and van Woesik, 2015). Understanding how coral taxa differ in their susceptibility to disturbances is integral to predicting how coral communities may change in a warmer world (Côté and Darling, 2010).

Highly disturbed reefs provide an opportunity to investigate the potential impacts of climate change, both on coral populations and communities. The Keppel Islands, located in the Southern inshore Great Barrier Reef (GBR), lie approximately 30 km from the mouth of the Fitzroy River, which drains the largest GBR catchment (Furnas 2003). These rocky continental islands and outcrops, sited in a shallow sandy bay, experience a 4 m tidal range that generates strong currents and high turbidity (Furnas, 2003; van Woesik and Done, 1997; Water Technology Water Coastal Environmental Consultants, 2013). Reefs in this region are considered ‘highly disturbed’ and have been impacted by six major flooding events, four cyclones, four major storms, and six coral-bleaching events driven by marine heat waves over the past 30 years (**Figure 1**) (Diaz-Pulido et al., 2009a; Jones and Berkelmans, 2014; Thompson et al., 2021; van Woesik et al., 1995). Bleaching in 1998, 2002 and 2006 was severe (greater than 60% of coral cover bleaching)(Berkelmans et al., 2004; Diaz-Pulido et al., 2009b), while bleaching in 2016 and 2017 was comparatively mild (Kennedy et al., 2018; Thompson et al., 2021). Despite frequent disturbances, coral communities in the Keppel Islands continue to be dominated by disturbance-sensitive branching *Acropora* species (Thompson et al., 2021). Rapid recovery of *Acropora* via asexual fragmentation and re-growth of remnant tissue, combined with rapid *Acropora* growth rates documented in this region (Diaz-Pulido et al., 2009a), may, at least in part, explain the ability of *Acropora* colonies to rapidly recover from disturbances and maintain their dominance in this region (Diaz-Pulido et al., 2009b; Thompson et al., 2021). Local adaptation to these disturbances is also supported by genetic data indicating that the Keppel Islands *Acropora millepora* population receives little genetic input via sexual recruitment from other GBR populations (van Oppen et al., 2015). Self-seeding via sexual recruitment is also likely to contribute to reef recovery following disturbances, however in recent years recruitment of Acroporidae in the region has been historically low (Davidson et al. 2019).

**Figure 1.**
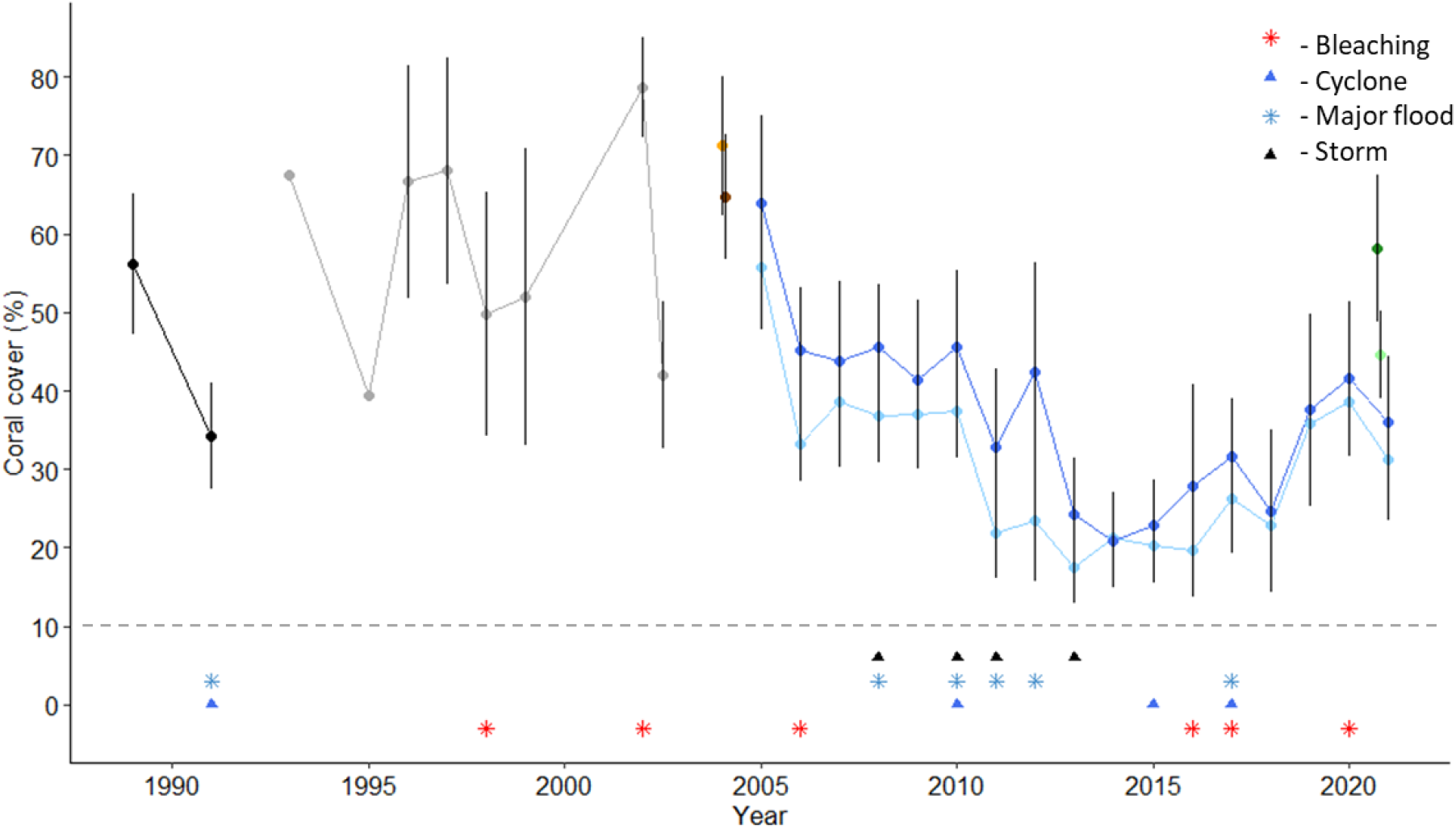
Historic cover of scleractinian corals in the Keppel Islands from 1989 to 2021. Disturbances: Red astericks = bleaching, black triangles = storms, blue triangles = cyclones, blue astericks = floods. Data sources: Black = van Woesik 1991 (van Woesik, 1991), Grey = Queensland Department of Environment and Heritage (unpublished data), Orange (2m) and Brown (5m) = Marine Tropical Sciences Research Facility (MTSRF)(Sweatman et al., 2007), Light blue (2m) and Dark blue (5m) = Great Barrier Reef Marine Park Authority Marine Monitoring Program (Thompson et al., 2021), Light green (1-4m) and Dark green (4-7m) = this study.

Regional-scale bleaching events provide an opportunity to explore how spatial heterogeneity in environmental conditions, including water flow and heat stress, contribute to variation in bleaching and recovery and can improve our ability to predict how coral communities may fare in a warmer climate. In this study, we analysed a bleaching event that impacted much of the GBR in early 2020 (Hughes and Pratchett, 2020), to (i) characterise the heat stress experienced across Keppel Island sites using remotely sensed and modelled seawater temperature and ocean current data, compared with a historical climatology, (ii) track coral bleaching and survival amongst taxa and across depths at six reef sites, (iii) quantify site-specific bleaching-induced mortality in *Acropora millepora*, and (iv) investigate the role of heat stress and flow rate in driving bleaching and recovery. Results of this study provide insights into the current resilience of coral communities to heat exposure and bleaching in this highly disturbed region, and highlights the likelihood of this region maintaining its current coral community composition into the near future.

## Methods

### Regional-scale climatology

To provide context for the thermal history of the Keppel Islands, long-term (150 year) mean monthly sea surface temperatures (SSTs) from Jan 1870 to Jan 2021 were obtained from the Met Office Hadley Centre reconstructed sea surface temperature (HadISST) record at a course gridded 1° by 1° spatial resolution (Rayner, 2003). The data were obtained from a centrally located point within the grid-cell that encompasses the greatest coverage of the Keppel Islands (23°30’0”S, 151°30’0”E). A climatological monthly mean temperature (CMM) was constructed from an 80-year reference window of 1870 – 1950 and monthly thermal anomalies (SST_A_) from 1870-2021 were calculated as the difference between the monthly temperature (MT) and the climatological monthly mean (CMM) (Equation 1).

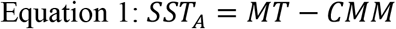

Trends in monthly SST, monthly SST_A_, and annual SST minima and maxima were visualised in R (R Core Team 2020) using the package ‘ggplot2’ (Wickham, 2016). A linear model was used to evaluate the rate of change of each SST metric in base R (R Core Team 2020). To examine the frequency of thermal anomalies in the Keppels, and to assess their temporal distribution over the last 150 years, the detrended and centred monthly SST_A_ record was deconstructed into time-frequency space using spectral analysis. A Morlet univariate wavelet transform model was generated using the ‘biwavelet’ package (Gouhier et al. 2018) in R (R Core Team 2020), and the frequencies were confirmed by evaluating spectral densities.

### Site-specific climatology and degree heating week calculations

Daily modelled seawater temperature (°C), eastward current (m s^-1^) and northward current (m s^-1^) data were extracted for each of the study sites from the GBR1 Hydro V2 eReefs model^1^ for the entire available temporal range (1 December 2014 to 12 February 2021). Modelled temperature and current data were extracted at a depth of 2.35 m and 5.35 m for the shallow and deep sites, respectively. A climatological weekly mean temperature (CWM) for each site and depth was calculated from 2015 – 2019 and a modified degree heating week (DHW) metric was calculated from 1 Jan to 31 April 2020 as the sum of the difference in the weekly maximum daily temperature (WMT) and the climatological weekly mean temperature (CWM), when greater than 1, divided by 7 (Equation 1). The modified DHW was used to characterise the level of heat exposure experienced by the corals at each site during the height of the heat wave and represents a conservative estimate of heat exposure given the recent baselines used in these calculations.

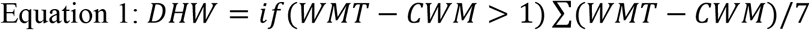

To investigate the role of flow rate in bleaching and recovery, the absolute values for eastward and northward (m s^-1^) currents were calculated for each site, to account for water flow in both positive (eastern or northern) and negative (western or southern) directions. Then, the absolute values of the median and maximum daily currents were averaged from January – June 2020, during the height of the thermal stress and recovery periods.

### Field surveys of coral bleaching and benthic cover

To document the severity of coral bleaching during the 2020 marine heat wave, surveys were completed at five reefs in the Keppel Island region in early April 2020 following reports of widespread and severe bleaching (**Supplementary Table 1**). Reefs were haphazardly selected and were chosen with no prior knowledge of the severity of bleaching. Photo-transect surveys were completed at both a shallower upper reef slope (∼1 – 4 m below the lowest astronomical tide (LAT)) and a deeper lower reef slope (4 – 7 m below LAT) site at each reef, except at Pumpkin Island and Shelving Bay (Great Keppel Island) where surveys were only completed on the shallow reef slope due to time constraints and the absence of a deep reef slope, respectively. Surveys were repeated at these and an additional three sites in June and October 2020 (also selected haphazardly to represent a larger number of reefs, **Figure 2E**) to document the impact of, and recovery from, bleaching amongst sites and taxa (**Supplementary Table 1**). Photo transect and photo sampling methods closely followed those used by the Australian Institute of Marine Science (AIMS) Long-term Monitoring Program (Jonker et al., 2008) and are briefly outlined below.

**Figure 2.**
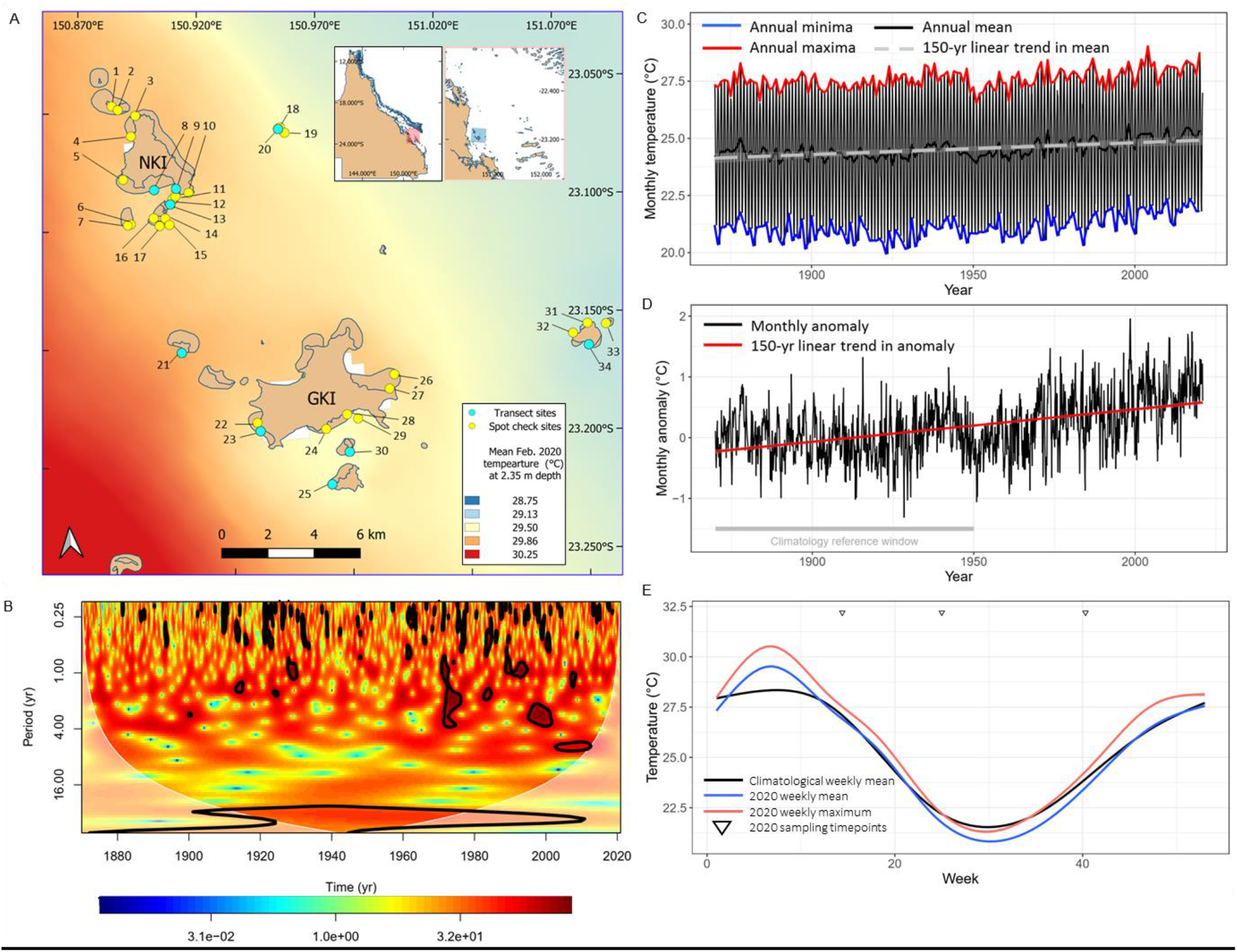
(A) Map of the Keppel Islands region with field sites identified. Numbered sites are defined in **Supplementary Table 1**. Map background colour represents the mean temperature in February 2020 at the height of the heat wave, at 2.35 m depth, as modelled by eReefs (www.ereefs.aims.gov.au). (B) Morlet wavelet model output indicating the periodicity of sea-surface temperature anomalies in the Keppel Islands from 1870 to 2020 (from D). Colour bar represents the power spectrum, indicating the strength of the signal in time-frequency space. All regions within black contour lines represent 95% confidence limits of statistically significant periodicities (*P* ≤ 0.05). The white line indicates the ‘cone of influence’ and periodicities identified outside this region (shaded area) should be approached cautiously. (C) Monthly sea surface temperatures in the Keppel Islands from 1870 to 2020 with annual maxima and minima. Mean and 150-yr trend lines are overlaid. (D) Monthly sea surface temperature anomalies in the Keppel Islands from 1870 to 2020, with the linear trend overlaid, and the reference window for anomaly calculations identified. (E) The weekly recent climatological mean, with the 2020 weekly mean and maximum overlaid for the Keppel Islands.

At each site, five 20 m transects were haphazardly laid along the deep and shallow reef slopes along the reef contour. The start of each transect was separated from the end of the previous transect by at least 5 m. On one side of the transect, photos of the benthos were taken at 50 cm intervals, ensuring that photos did not overlap. The camera lens was held parallel to the reef substrate and the camera height above the substrate was maintained at ∼ 1 m to ensure consistency in the surface area included in each photograph (approximately 25 cm^2^). The number of photographs recorded per transect averaged 31 ± 10 (SD).

### Photographic analyses

Estimates of the benthic community composition were derived from point-count analyses. Ten points were overlaid in an even distribution on each photograph and the benthic organism under each point was identified to the lowest possible taxonomic level, with most Scleractinia (stony corals), Alcyonacea (soft corals), and macroalgae identified to genus. For points identified as stony corals (hereafter ‘corals’), the level of bleaching was recorded as either not bleached, pale or fully bleached, consistent with methods described in Jonker et al. 2008. Alcyonaceans were examined for bleaching, but none showed signs and were thus excluded from the analysis. Coral points scored as not bleached appeared normally pigmented and showed no signs of paling (corresponding to a score of 5 or 6 on the Coral Watch Coral Health Chart, although the Chart was not used to score points in this study (Siebeck et al., 2006)). Corals scored as pale were fluorescent in colour or lighter in colour than normal for that taxon, but not bright white (corresponding to a score of 2-4 on the Coral Health Chart). Bleached corals appeared bright white, corresponding to a score of 1 on the Coral Health Chart.

To examine spatial variation in bleaching more widely, spot checks were completed at an additional 25 sites in June 2020 (**Supplementary Table 1)**. Representative photos of coral communities at these spot check locations were taken on snorkel. Images were taken parallel to the reef benthos and from 1 – 3 m above the benthos. For each site, the benthic composition and bleaching severity and prevalence were determined from ten photos randomly chosen from those taken at each location, as described above for photo transects (Jonker et al., 2008). Scleractinian points scored as bleached and pale were pooled and collectively referred to as “bleached” in all statistical analyses.

### Statistical analyses of bleaching

Temporal trends in bleaching were investigated by modelling the total numbers of bleached and unbleached scleractinian points per transect against survey month and depth and their interaction (fixed factors) using a generalised linear mixed effects model (GLMM) with transect nested in reef included as a random effect (random intercept) and using a beta-binomial distribution and a logit link function. Models were run using the ‘glmmTMB’ package (Brooks et al., 2017). Model assumptions including homogenous variance and normally distributed residuals were verified using the ‘DHARMa’ package (Hartig, 2021). For significant models, pairwise comparisons were undertaken by calculating least-squares mean with a Tukey’s adjustment using the ‘emmeans’ package (Lenth, 2021).

Differential susceptibility to bleaching across coral genera was investigated by modelling the total numbers of points scored as bleached or not bleached at the height of bleaching in April 2020 against genus (fixed factor) using a GLMM. Replicate transects nested in depth and reef was included as a random effect (random intercept). Data were modelled using a binomial distribution and a logit link function. Only those genera representing at least an average of 5% of coral cover across transects were included in this analysis (*Acropora, Pocillopora, Montipora* and *Porites). Pocillopora* corals were then further excluded from the analysis as nearly 100% of points were bleached. Model assumptions of homogeneity of variances and normally distributed residuals were verified using the package ‘DHARMa’ (Hartig, 2021).

To determine whether differential taxonomic susceptibility to bleaching was consistent across depths, the total numbers of points scored as bleached and not bleached were modelled against the interaction between genus and depth (fixed factor) using a GLMM. Transect nested in reef nested in depth was included as a random effect in this model with a beta-binomial distribution and logit link function. Only the genera *Acropora* and *Montipora* were included in this model as these were the only genera for which sufficient data were available from both depths. Model assumptions were verified and post-hoc pairwise comparisons completed as described above.

A GLMM was also fitted to investigate whether the cover of scleractinian corals declined over the six months following bleaching. Survey month and depth and their interaction were included as fixed factors and transects nested in reef were included as a random factor, to account for variation among reefs. Data were modelled with a beta-binomial distribution and logit link function, as described above. All statistical analyses were completed in R (R Core Team, 2020) and data visualisations completed using the R package ‘ggplot2’ (Wickham, 2016).

### Fate-tracked colonies

Estimates of whole-colony mortality following bleaching were determined from *Acropora millepora* colonies that were fate-tracked at four reefs during the bleaching event (18 – 21 April 2020) and resurveyed in October 2020. Adult colonies (9.5 – 73 cm in diameter, average diameter = 30 cm) were haphazardly chosen to represent the range of colony responses to heat stress (from severely bleached to non-bleached), although given the severity of bleaching in the region, few colonies (∼11% colonies) remained unbleached. Approximately 100 colonies were tagged on the shallow reef flat (∼ 1 – 2 m below LAT) at each of Pumpkin Island, North Keppel Island and Halfway Island, while 50 colonies were tagged at Great Keppel Island, for a total of 350 tagged colonies. Colonies were located up-slope of photo transect sites at these same locations, as that is the primary habitat of *Acropora millepora* in the Keppels. In October 2020, colonies that could be found were resurveyed and recorded as dead or alive.

### Relationships between bleaching recovery and environmental variables

The relationships between environmental conditions (heat and flow metrics) and both bleaching (in April 2020) and recovery from bleaching (between April and June 2020) were examined by modelling the maximum daily temperature during the 2020 heat wave, the DHW values, and the average flow rates against (i) bleached *Acropora* cover and (ii) the change in the proportion of bleached *Acropora* cover between April and July across the photo-transect sites that were surveyed in both months (n = 5 reefs at each of two depths). To explore the relationships between the environmental conditions (heat and flow) that persisted through to June 2020 and bleaching, the same environmental metrics were modelled against the proportion of *Acropora* cover bleached in June across all transect and spot-check sites (n = 42).

The data, as described above, were modelled with a linear model using generalised least squares fit by restricted maximum likelihood (REML). Because the sites were clustered in space (**Figure 2E**), a Moran’s I test for distance-based autocorrelation was calculated for each model using the DHARMa package (Hartig, 2021), and where spatial autocorrelation was significant (at *p* < 0.05), a spatial dependence correlation structure was applied to the models. The structures tested included an exponential, a gaussian, a rational quadratic, and a spherical structure, and the best model was selected based on the lowest Akaike Information Criteria (AIC) estimates and verified using a semivariogram. Models were run using the ‘gls’ function from the R package ‘nlme’ package (Pinheiro et al., 2021).

## Results

### Long-term temporal trends in heat stress in the Keppel Island region

Average sea-surface temperatures in the Keppel Islands have increased 0.8 °C over the past 150 years (**Figure 2A and B**). The rate and frequency of anomalously warm months is also increasing, with evidence of significant frequent (1 – 4 year) anomalies since about the 1970s (**Figure 2D**). The increase in average temperatures is driven by increases in both annual temperature minima and maxima (**Figure 2B**), with annual minima increasing at a faster rate (0.9 °C 150 yr^-1^) than annual maxima (0.7 °C 150 yr^-1^). Since February 2014, 90% of the monthly temperature anomalies in the Keppel Islands have been warmer than the climatological average (**Figure 2B**).

During the 2020 marine heatwave, average weekly seawater temperatures in the Keppel Islands started to exceed the recent (2015 – 2019) climatological mean temperature (CMT) in mid-January 2020 and continued to exceed the CMT for 8 weeks until mid-March (**Figure 2C**). Maximum water temperatures in the Keppel Islands rose sharpy and peaked in mid-February at 32.0 °C at Miall Island, 3.5 °C degrees above the climatological mean (**Figure 2C**). Cumulative heat stress, measured as DHWs from 1 January to 31 April 2020, ranged between 3.75 and 7.95 and was highest inshore of the northern islands group (North Keppel, Conical, Corroboree, Pumpkin and Sloping Islands and Square Rocks) as well as inshore of Miall Island, which lies to the north of the southern island group (**Figure 2E**). Cumulative heat stress was lowest at the offshore sites: Barren Island, The Child, Bald and Outer Rocks (**Figure 2E**). Maximum daily seawater temperatures between 1 January and 31 April 2020 varied between 26.1 and 32.0 °C. The Summer/Autumn 2020 marine heat wave was followed by a cooler than average Winter/Spring in the Keppel Islands, relative to the recent climatology (2015-2020). Water temperatures were cooler than the climatological mean by as much as 1.2 °C (**Figure 2D**).

### Patterns in bleaching severity in April 2020

In early April 2020, coral bleaching was widespread and severe at all sites surveyed (**Figure 3, Figure 4**) and at other sites throughout the Keppel Islands from which reports of bleaching were available (Hughes and Pratchett, 2020; North Keppel Island Environmental Education Centre, 2020). Surveys revealed that over 75% of live coral was bleached at all sites, with ≥90% of live coral cover bleached at seven of the ten sites (**Figure 4**). Bleaching was most prevalent on the shallow reef at Barren Island (98.5%), and lowest on the shallow reef at Halfway Island (75%) (**Figure 4**). Very recent coral mortality, evident as bright white coral skeleton denude of coral tissue and fouling organisms, was recorded from all sites but represented less than 3% cover. No bleaching was recorded in any other taxonomic group (i.e. soft corals, anemones, clams) although these taxa are not common on the reefs surveyed. The prevalence of bleaching on deeper (4 – 7 m) reefs was as high, or higher than the shallow reefs (1 – 4 m) (GLMM: *z =* −4.196, p < 0.001: **Table 1**). On average 92.3 ± 1.2% of coral cover was bleached at deep sites compared with 89.7 ± 1.7 % at shallow sites (**Figure 5A**).

**Table 1:** Generalized linear mixed effects model summary statistics predicting the probability of a coral being bleached based on sampling month and depth. Significant *p*-values are indicated in bold.

| Random effect (intercept) | Variance | Std.Dev |  |  |
| --- | --- | --- | --- | --- |
| Transect:Reef | 1.358e-09 | 3.685e-05 |  |  |
| Reef | 1.388e-01 | 0. 3.726e-01 |  |  |
| Fixed effects | Estimate Std. | SE | Z value | Pr(>\|z\|) |
| Intercept (April 2020-Deep) | 0.5246 | 0.2033 | 2.580 | **0.00987** |
| Shallow | -0.7034 | 0.1676 | -4.196 | **2.72e-05** |
| June 2020 | -2.1268 | 0.2070 | -10.276 | **< 2e-16** |
| October 2020 | -4.0910 | 0.2999 | -10.276 | **< 2e-16** |
| Shallow-June 2020 | 1.0647 | 0.2558 | 4.163 | **3.14e-05** |
| Shallow-October 2020 | 0.3738 | 0.3729 | 1.002 | 0.31623 |

**Figure 3.**
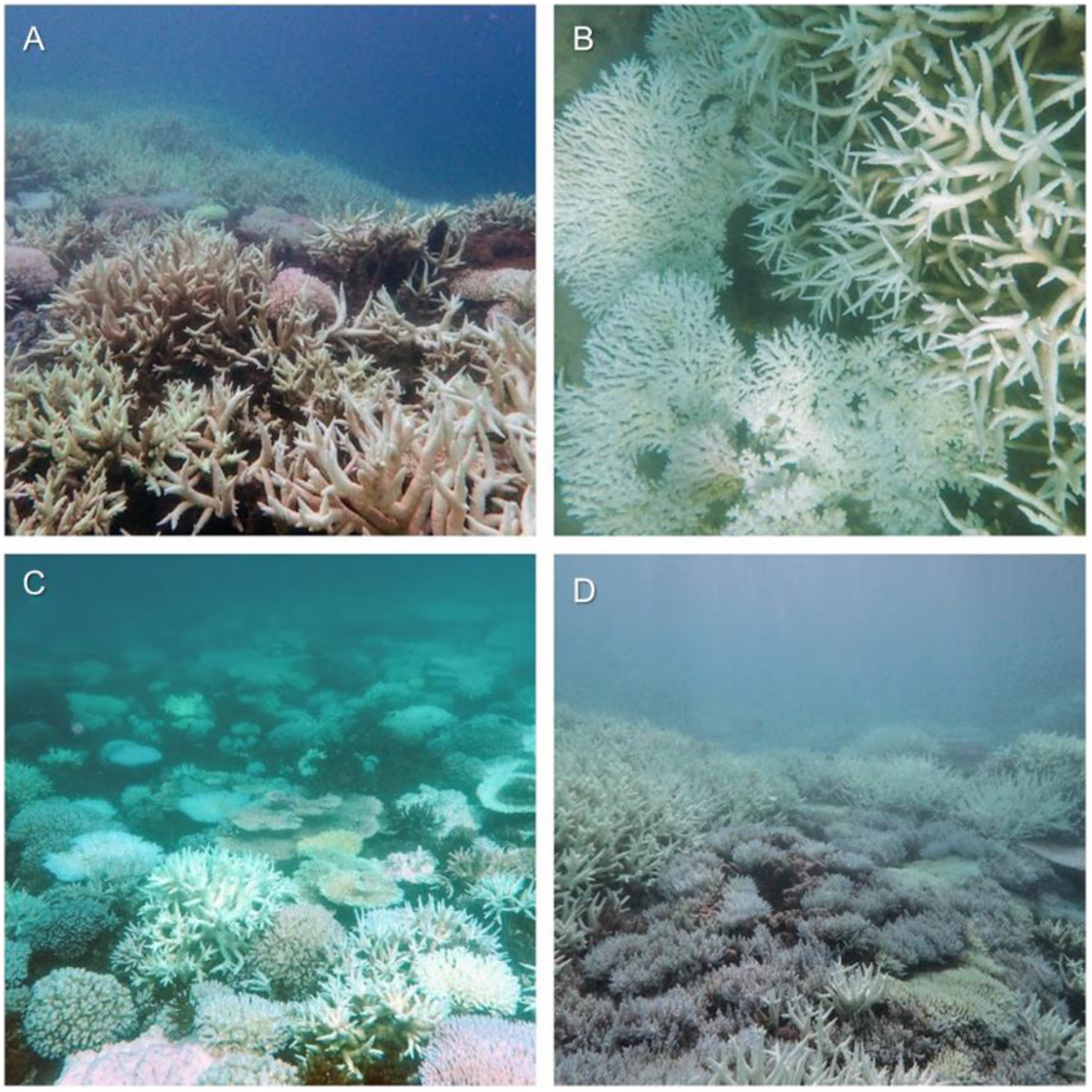
Examples of bleached corals in April 2020, at (A) Outer Rock, (B) Mazie Bay, North Keppel Island, (C) Halfway Island and (D) Barren Island. Photo credits: (A and B) Darren Brighton, (C and D) Scott Gardner.

**Figure 4:**
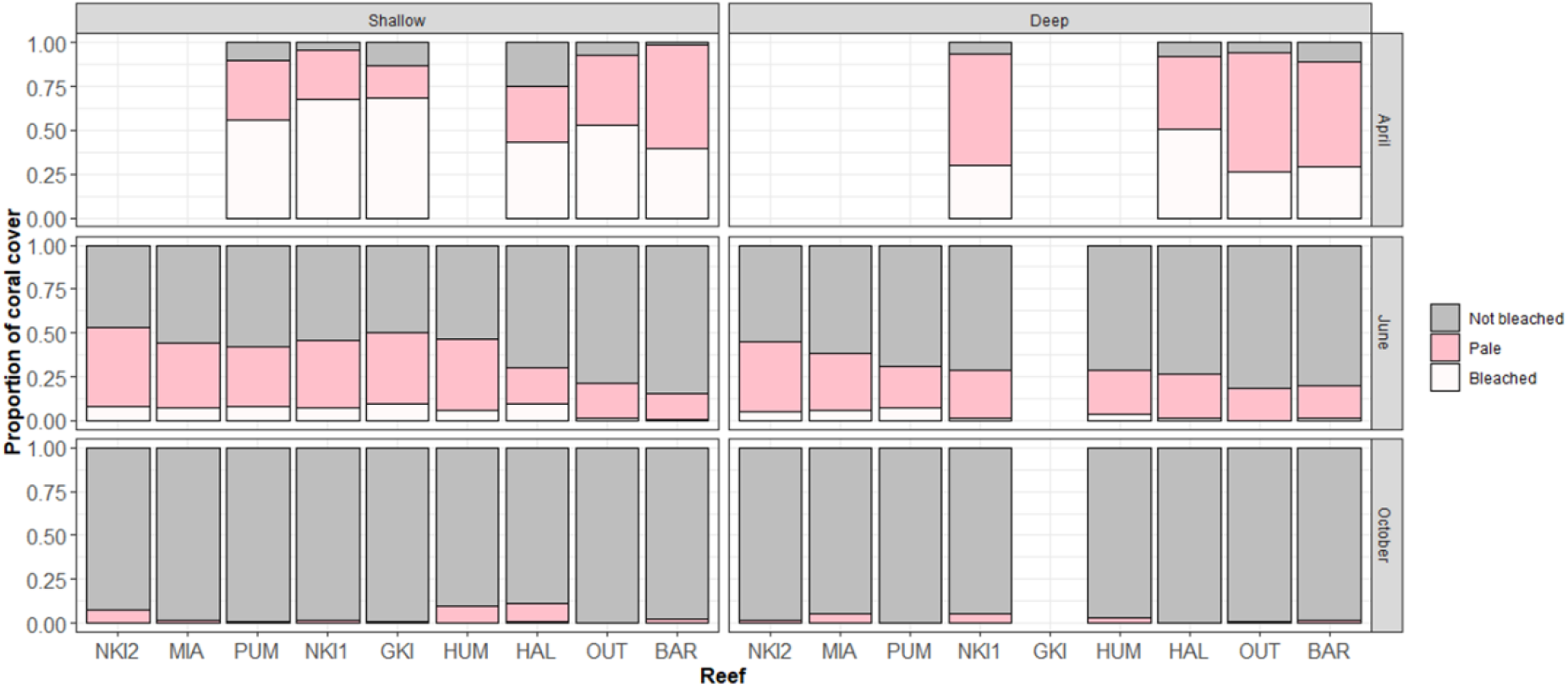
Proportion of coral cover bright white (bleached), pale, or normal in pigmentation at photo-transect sites in April, June and October 2020. Site abbreviations are defined in **Supplementary Table 1**.

**Figure 5.**
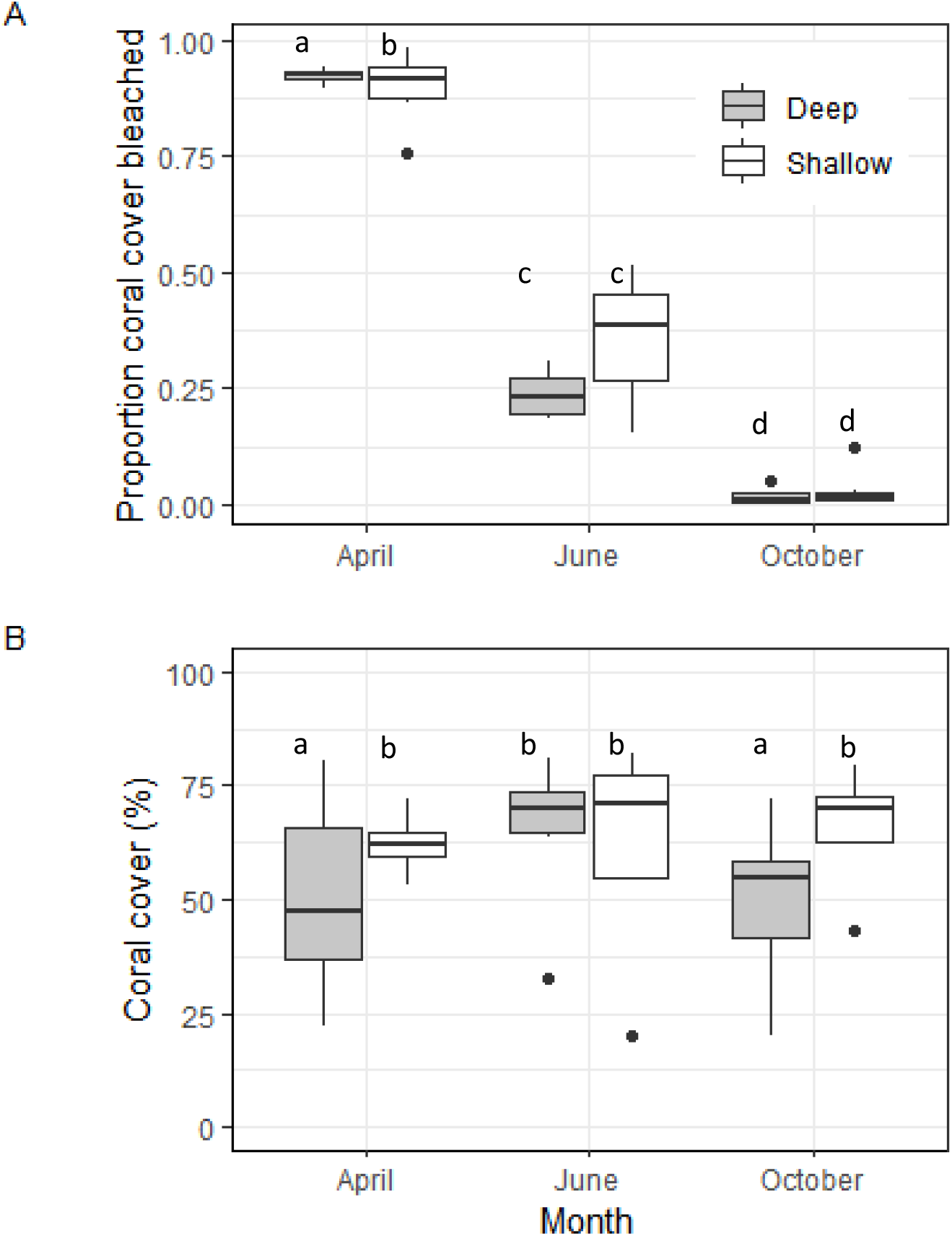
Temporal trends in (A) the proportion of corals bleached and (B) the percentage cover of scleractinian corals at Keppel Island reefs in April, June and October 2020, across two survey depths.

Repeat surveys of those sites surveyed in April 2020, combined with additional sites surveyed using photo transect and spot check methods, revealed the persistence of bleaching in June at all but one site (East Clam Bay, GKI, **Supplementary Table 1**). Recovery was more rapid at deep sites compared with the shallow sites (GLMM: Shallow – June 2020: *z* = 4.163, p = <0.001: **Table 1**). By June 2020, the proportion of corals bleached was similar at deep (23.8 ± 2.3 % cover) and shallow depths (35.6 ± 2.9 % cover; pairwise comparison: p = 0.08; **Figure 5A**). By October 2020, bleached corals were rare and occurred at only the shallow reef at Halfway Island. Pale corals were recorded at all sites, but these generally constituted less than 10% of coral cover (**Figure 4**). The percent cover of bleached coral cover was similarly low at both shallow (3.4 ± 1 % cover) and deep sites (0.19 ± 0.6 % cover pairwise comparison: p = 0.34; **Figure 5A**).

### Impact of bleaching on coral survival and cover

Despite the severity of bleaching, survival of *A. millepora* colonies was high (**Figure 6**). No whole-colony mortality occurred at North Keppel Island, while whole-colony morality was highest at Pumpkin Island (17% of colonies)(**Figure 6**). Consistent with these results, average cover of corals in October 2020, six months after bleaching, did not differ from that initially recorded in April 2020 at both shallow and deep sites (Shallow sites: April cover = 50.4 ± 2.5%, October cover = 49.7 ± 1.9%, pairwise comparison: p = 0.93; Deep sites: April cover = 62.3 ± 2%, October cover =65.5 ± 2.5%, Pairwise comparisons p = 0.87) (**Figure 5B, Table 2**).

**Table 2.** Generalized linear mixed effects model summary statistics predicting the proportional cover of scleractinian corals based on sampling month and depth. Significant *p*-values are indicated in bold.

| Random effect (intercept) | Variance | Std.Dev |  |  |
| --- | --- | --- | --- | --- |
| Transect:Reef | 1.301e-09 | 3.607e-05 |  |  |
| Reef | 2.343e-01 | 4.841e-01 |  |  |
| Fixed effects | Estimate Std. | SE | Z value | Pr(>\|z\|) |
| Intercept (April 2020 - Deep) | 0.72877 | 0.26833 | 2.716 | 0.0066* |
| Shallow | -0.69702 | 0.22479 | -3.101 | **0.0019** |
| June 2020 | -0.05672 | 0.23682 | -0.240 | 0.81073 |
| October 2020 | 0.11750 | 0.23858 | 0.492 | 0.8107 |
| June 2020 - Shallow | 0.64287 | 0.30668 | 2.096 | 0.0360 |
| October 2020 - Shallow | 0.18397 | 0.30508 | -0.603 | 0.5465 |

**Figure 6:**
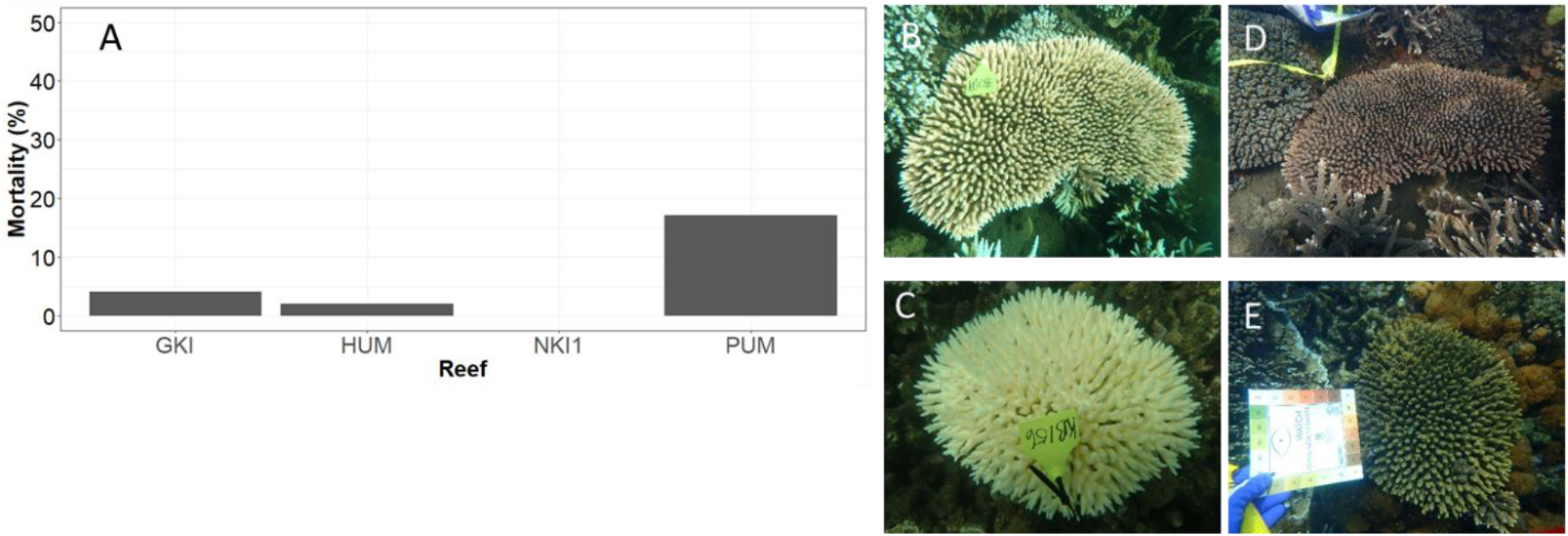
(A) Mortality of fate-tracked colonies of *Acropora millepora* at four sites in the Keppel Islands between April and October 2020. Photos demonstrate examples of bleached colonies in April (B and C) that had recovered by October 2020 (D and E). Photo credits: (B and C) D. Brighton, (D and E) C. Randall.

### Differential susceptibility of scleractinian taxa

Keppel Island coral communities were dominated by the genus *Acropora*, particularly the branching species *A. muricata* and *A. intermedia*. In April 2020, on average 48.5 ± 6% of coral cover was comprised of *Acropora* colonies (**Figure 7A**). Only three other coral genera, *Pocillopora, Porites* and *Montipora*, were present at greater than an average of 5% cover per transect (**Figure 7A**). These common coral taxa were also the most susceptible to bleaching (**Figure 7B**) with 96.6 ± 3.4% of *Pocillopora* colonies bleached. An average of 94.6 ± 0.9% of *Acropora*, 66 ±12.9 % of *Porites* and 43.6 ± 5.8 % of *Montipora* points were recorded as bleached (**Figure 7B**). *Acropora* and *Porites* corals were equally susceptible to bleaching but *Porites* colonies exhibited a greater range of bleaching responses (GLMM: *z* = −0.83, p = 0.407: **Table 3**). Both *Acropora* and *Porites* corals were more susceptible to bleaching than *Montipora* colonies (Pairwise comparison: p < 0.001; **Table 3)**. The susceptibility of the two most common coral genera, *Acropora* and *Montipora*, did not vary between depths, with *Acropora* corals being more susceptible than *Montipora* corals at both depths (Pairwise comparison: p = 0.001; **Table 4**).

**Table 3.**
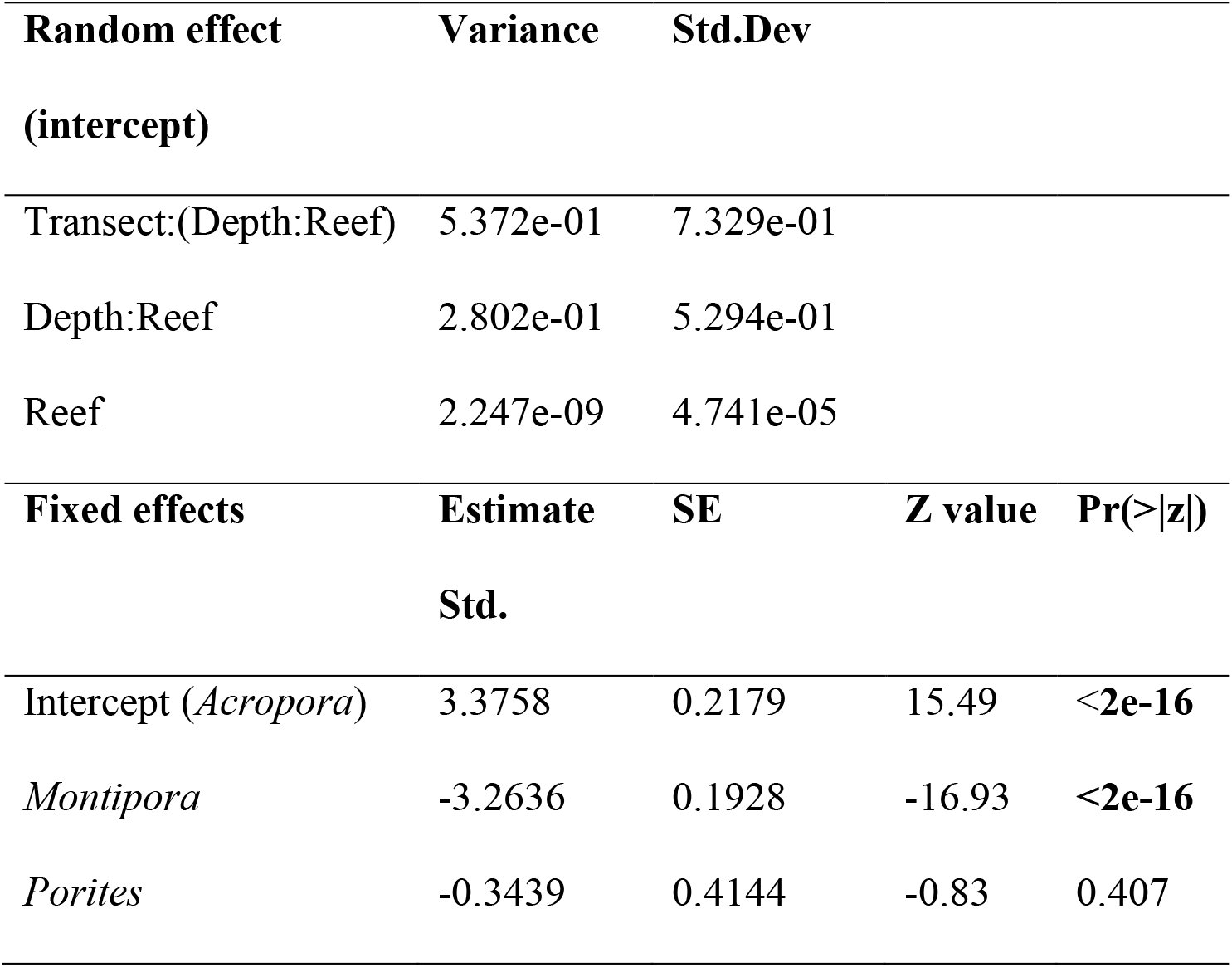
Generalized linear mixed effects model summary statistics predicting the probability of a coral being bleached based on its taxonomic identity (*Acropora, Montipora, Porites* only). Significant *p*-values are indicated in bold.

**Table 4:**
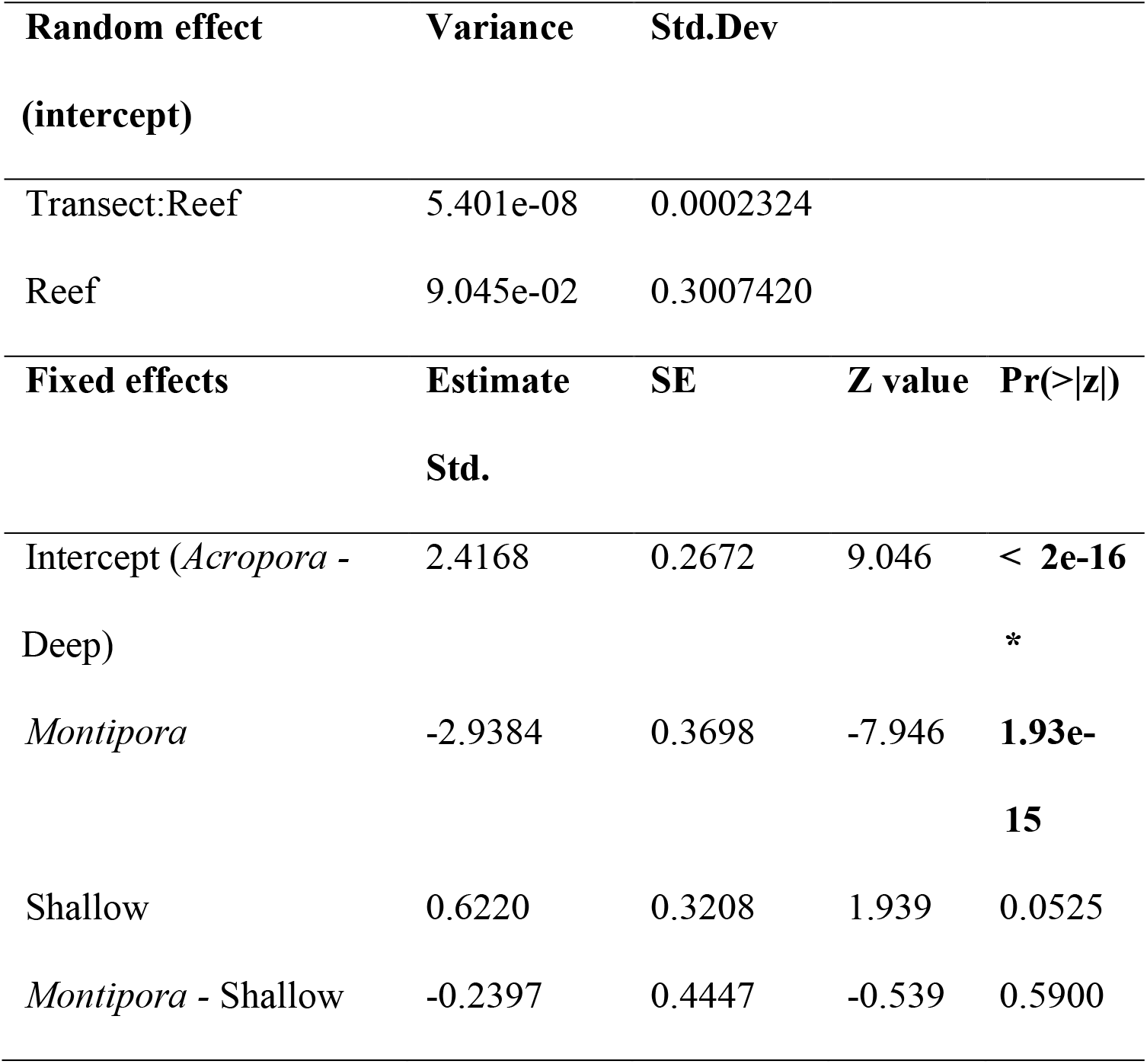
Generalized linear mixed effects model summary statistics predicting the probability of a coral being bleached based on its taxonomic identity and depth (*Acropora* and *Montipora* only). Significant *p*-values are indicated in bold.

**Figure 7.**
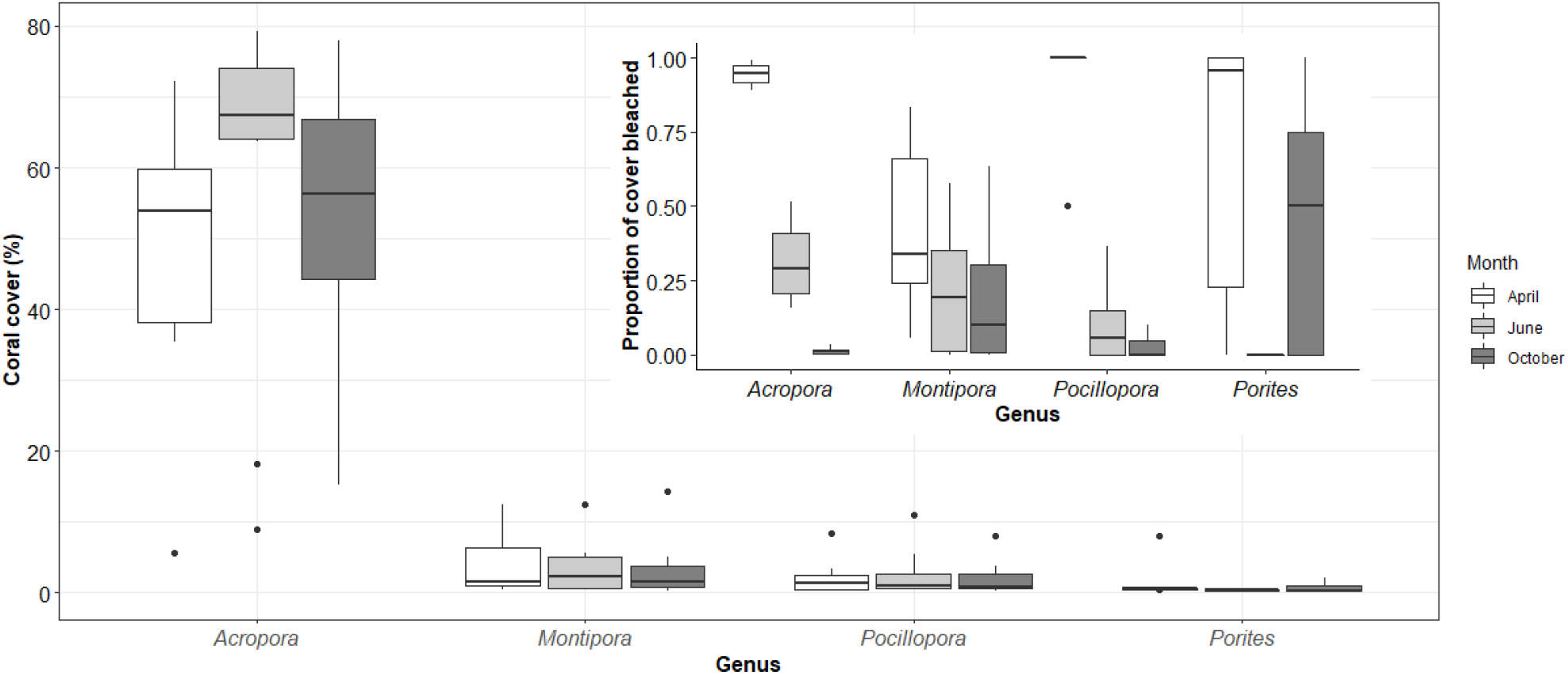
Average cover (%) of scleractinian coral genera across Keppel Island reefs (main plot), and the average proportion of cover scored as bleached (inset), in April, June and October 2020.

### Relationship between environmental metrics and recovery from bleaching

Bleaching severity in April 2020 did not correspond with metrics of heat-exposure or flow rates, likely due to high bleaching (75 – 98%) at all sites as a consequence of exceeding heat-exposure thresholds (**Figure 2C, Figure 8**). All sites, including both shallow and deep reefs, experienced four or more degree heating weeks and at least a maximum daily temperature of 29.5 °C from January to April 2020. The persistence of bleaching in June, however, was driven by accumulated heat-exposure and for every additional 1 DHW of exposure, an additional ∼10% of corals remained bleached (**Figure 8; Supplementary Table 2**). Similarly, for every 1 °C increase in the maximum daily temperature experienced, an additional ∼30% of corals remained bleached (**Figure 8**; **Supplementary Table 2**). Heat-exposure in the lead up to bleaching was, therefore, a strong predictor of persistent bleaching in June 2020. Recovery from bleaching was slower at inshore reefs where heat-exposure was highest. Persistent bleaching in June 2020 was also often higher at shallow sites compared to deep sites having experienced the same exposure to accumulated heat (**Figure 8**).

**Figure 8:**
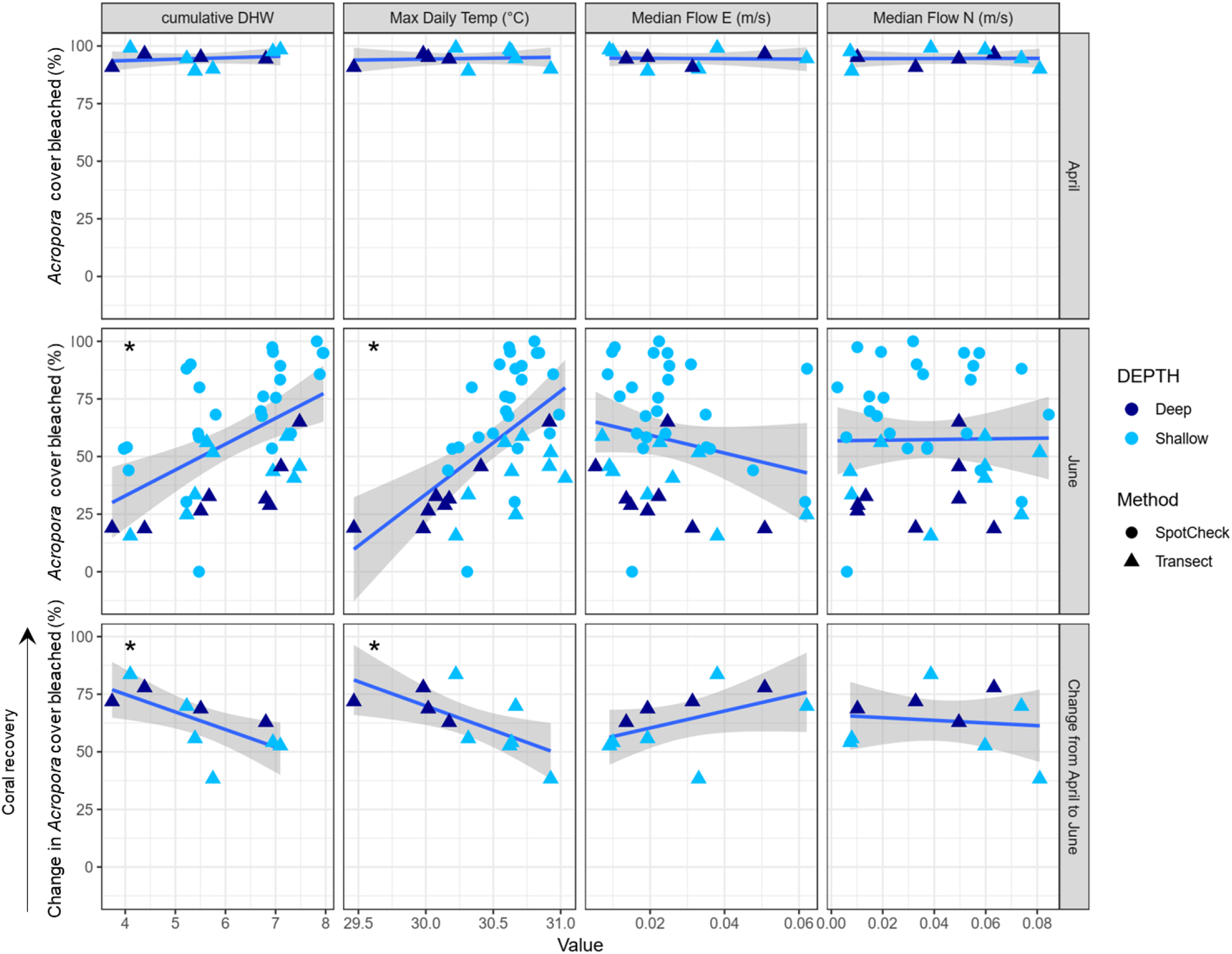
Relationships between the proportion of *Acropora* corals bleached in April and June 2020 (top and middle rows, respectively) and a suite of environmental metrics (columns; cumulative degree heating weeks (DHW), maximum daily temperature, and median flow rates in east – west and north – south directions). The bottom row indicates the relationship between the change in bleaching from April to June, as a proxy for recovery, and the environmental metrics (columns). Asterisks indicate statistically significant trends (**Supplementary Table 1**) and grey shading represents the 95% confidence intervals around the models.

Neither bleaching severity in April and June, nor the recovery from bleaching, was significantly predicted by median flow rates, despite a tendency toward higher recovery and lower bleaching at higher-flow sites. Flow rates were significantly correlated with cumulative DHW and higher heat exposure was recorded at sites with lower east-west flow rates (F = 21.03, *p* < 0.001, *r*^*2*^ = 0.328) and where rates of northerly water flow were highest (F = 3.377, p = 0.015, r^2^ =0.116; **Supplementary Figure 1**).

## Discussion

During early 2020, over 90 % of shallow-water corals in the Keppel Islands bleached (**Figure 4**). This represents a much higher severity of bleaching than recorded offshore at One Tree Island (∼ 47% of coral cover) two months earlier (Nolan et al., 2021), which reflects higher overall temperatures recorded at the Keppel Islands (>30 °C compared to < 30 °C at One Tree Island) and potentially greater accumulation of heat exposure over these two months at the Keppel Islands. The severity of bleaching in the dominant *Acropora* genus was high across sites, reflecting high heat stress across the entire region and therefore did not correlate with site-specific heat-stress metrics, in contrast to many historic studies (Berkelmans, 2001; Donovan et al., 2021; Glynn, 1993; Harrison et al., 2019; Heron et al., 2016; Hughes et al., 2017b, 2019b). These results add to the growing evidence that coral bleaching thresholds are being exceeded across entire regions during marine heat waves and emphasizes the decoupling of bleaching thresholds and bleaching severity. The occurrence of more frequent and severe marine heat waves under projected climate warming scenarios (Cai et al., 2014; Hoegh-Guldberg, 1999; Hughes et al., 2017a; Timmermann et al., 1999), coupled with the accumulation of latent impacts of heat stress over multiple years, is likely to lead to a further decoupling of bleaching severity and the heat stress metrics that are currently in use (Hughes et al., 2019b; Neal et al., 2017). Metrics incorporating the accumulation of heat stress over multiple summers and accounting for the magnitude of winter reprieves, overlaid on background rates of temperature rise, are likely to be more useful in predicting bleaching severity in the future (Hughes et al., 2017b; Sully et al., 2019).

Severe bleaching events often result in high coral mortality (Brown, 1987; Glynn et al., 1998; Hughes et al., 2019a; Riegl, 2002; Stuart-Smith et al., 2018). Keppel Island corals however, exhibited high survival and stable coral cover following bleaching in early 2020 (**Figure 5, Figure 6**; Thompson et al., 2021). Yet, recovery of bleached colonies was faster on reefs with lower heat exposure. For every 1 additional DHW of exposure, ∼10% more corals remained bleached in June 2020. Similarly, for every 1 °C increase in the maximum daily temperature experienced, ∼30% more corals remained bleached in June 2020 (**Figure 8**; **Supplementary Figure 2**). High recovery across the region also may have been facilitated by cooler-than-average Autumn and Winter temperatures (**Figure 2C**). Nonetheless, results of this study indicate that even where temperatures exceed bleaching thresholds, small differences in heat exposure can affect recovery trajectories in the months following bleaching. Therefore, while the magnitude of heat exposure may be becoming less predictive of bleaching severity, it may remain useful for predicting recovery rates following future bleaching events. Severity of bleaching and short-term mortality remain the focus of many coral bleaching studies (Berkelmans et al., 2004; Harrison et al., 2019; Hughes et al., 2017b; P. J. Mumby et al., 2001). Results of this and other recent studies (Donovan et al., 2021; Stuart-Smith et al., 2018), highlight the importance of documenting coral survival or mortality over longer time frames in order improve our understanding of the contribution of environmental variables to heterogeneity in coral survival and the impacts of bleaching on coral community composition.

High water flow also can reduce bleaching and/or accelerate recovery, and the Keppel region is characterised by a large tidal range and strong currents (Furnas, 2003; Thompson et al., 2021; van Woesik and Done, 1997). High flow can rid corals of superoxide radicals through passive diffusion (Nakamura et al., 2005; Nakamura and van Woesik, 2001), and can increase water mixing to reduce temperatures. Water flow in this and other inshore regions is also typically correlated with the resuspension of particulate organic matter (Kleypas and Hopley, 1992), on which inshore corals are adept at feeding (Anthony, 2000). Heterotrophy can provide an important source of nutrition for corals in the absence of their symbiotic *Symbiodinium* populations and increase coral survival following bleaching (Grottoli and Palardy, 2006). Lower heat stress at sites with higher water flow in an east-west direction identified in this study, likely reflects greater mixing of deeper, cooler offshore waters with warmer shallower inshore waters (**Supplementary Figure 2**). In contrast, high flow rates in a north-south direction drove high temperature stress, most likely because north-south water flow resulted in the mixing of waters of similar depths, and the transport of warm water northward from the shallow Keppel Bay. High rates of water flow and increased heterotrophy by inshore corals is likely to be an important driver of the high tolerance of Keppel corals to bleaching. Finally, irradiance stress exacerbates bleaching (Downs et al., 2013; Lesser et al., 1990), and high turbidity can reduce this by reflecting light and shading corals (Cacciapaglia and van Woesik, 2016; Fisher et al., 2019; Morgan et al., 2017; Sully and van Woesik, 2020). Bleaching severity can be lower on turbid inshore reefs, such as the Keppel Islands, which may provide corals some protection from the effects of climate warming (Cacciapaglia and van Woesik, 2016; Fisher et al., 2019; Morgan et al., 2017; Sully and Woesik, 2020).

Consistent with previous studies, *Pocillopora* and *Acropora* were amongst the most susceptible genera to bleaching during the 2020 marine heatwave (Guest et al., 2012; Loya et al., 2001; Marshall and Baird, 2000; McClanahan et al., 2004)(**Figure 7B**). In contrast to previous studies *Porites* corals were also highly susceptible (Marshall and Baird, 2000; McClanahan et al., 2004). Resistance to bleaching in *Porites* is not universal (Guest et al., 2012; Mumby et al., 2001), and differential taxonomic susceptibility has proven to be temporally and spatially variable both within and among reef regions (Guest et al., 2012; Hughes et al., 2017b; Marshall and Baird, 2000; McClanahan et al., 2004). Sensitivity to bleaching does not always translate to high mortality (i.e. **Figure 6**) and both rates of mortality and sublethal impacts of bleaching are likely to be species-specific (Baird and Marshall, 2002; Cox, 2007; Johnston et al., 2020; Muir et al., 2017; Szmant and Gassman, 1990). Taken together, this highlights the need to document species-specific bleaching, mortality and sub-lethal responses, to provide a more comprehensive understanding of the impacts of bleaching for coral communities’ resilience (Côté and Darling, 2010; van Woesik et al., 2011).

The differential susceptibility of coral taxa to disturbance-mediated mortality can drive changes in the composition of coral communities (Aronson et al., 2002; Hughes, 1994; Johns et al., 2014; Loya et al., 2001; Pandolfi and Jackson, 2006), that may influence their vulnerability to future disturbances. The dominant *Acropora* corals in the Keppel Islands were highly susceptible to bleaching, yet survived (**Figure 5, Figure 6**,), indicating high resilience. The persistent dominance of *Acropora* over the past 30 years in this region, despite the occurrence of four cyclones, six bleaching events and six storms (**Figure 1**), to which the *Acropora* corals are highly sensitive (Fabricius et al., 2008; Loya et al., 2001; Madin, 2005; Marshall and Baird, 2000; van Woesik et al., 1995), supports this conclusion. High *Acropora* growth rates (Diaz-Pulido et al., 2009b) may support their ongoing dominance in the Keppel Islands. Low recruitment of non-*Acropora* species may also limit competition with other taxa on Keppel Island reefs, although *Acropora* recruitment is also lower than elsewhere (Davidson et al., 2019; Thompson et al., 2021). Disturbance sensitive coral taxa, including the *Acropora*, are forecast to be replaced by more resistant taxa on Indo-Pacific reefs under future warming scenarios (Côté and Darling, 2010; van Woesik et al., 2011). It is therefore unclear for how long current coral communities can retain the capacity to resist and recover from disturbances as these increase in frequency and intensity in a warming world (Cai et al., 2014; Côté and Darling, 2010; Harmelin-Vivien, 2021; Heron et al., 2016; Johns et al., 2014; Timmermann et al., 1999).

Rates of ocean warming vary in space (Dunstan et al., 2018; Heron et al., 2016; Randall and van Woesik, 2015) and within the southern GBR, long-term records indicate that sea surface temperatures have increased 0.8 °C since 1870 (**Figure 2A**). Corals here are experiencing warmer summers and winters, year on year, resulting in longer periods of summer-like temperatures and shorter winter reprieves, a pattern detected for many reefs worldwide (Heron et al., 2016). Minimum temperatures are also increasing faster than maximum temperatures, so stressful winter cold snaps that may have limited recovery or resulted in cold-water bleaching in the past (Hoegh-Guldberg et al., 2005; Hoegh-Guldberg and Fine, 2004; Howells et al., 2013), may be less problematic in the future (Schlegel et al., 2021). Shorter or less frequent winter reprieves from heat stress may also reduce recovery capacity of corals in the future. Faster recovery of coral at reefs experiencing lower heat exposure (**Figure 8**), combined with high survival of Keppel Island’s coral during the cooler than average Autumn and Winter that followed severe bleaching (**Figure 1C, Figure 6**), highlight the importance of cool reprieves for coral recovery in a warming world.

High spatial resolution satellite-derived sea-surface temperature models, such as that used here to calculate site-specific climatologies, are only available for a relatively recent window. For example, the GBR1 Hydro V2 eReefs model^2^ is only available from 2014 onwards. Heat-exposure metrics based on such recent baselines are therefore conservative and it is likely that DHW estimates would have predicted higher heat accumulation than was calculated here, had a longer-term baseline been available (**Figure 2D, Figure 8**). Despite the use of a recent baseline, this study still conservatively estimated the accumulation of 4 – 8 DHWs across all sites and depths, suggesting that heat exposure was indeed severe during the 2020 marine heat wave. Reefs exposed to more than 4 DHWs are expected to experience significant bleaching, while widespread severe bleaching and significant coral mortality is expected on reefs exposed to 8 or more DHWs (Skirving, 2020). Significant coral mortality was not recorded in this region, including on reefs experiencing almost 8 DHWs (7.95 DHWs), indicative of high resilience in this southern GBR and in contrast to the expectation of high mortality, given the naivety of the region to recent high heat exposure (Skirving, 2020).

Globally, the onset of coral bleaching is occurring at higher temperatures than previously, suggesting that directional selection for more thermally tolerant coral genotypes is underway (Sully et al., 2019). The high survival following severe heat exposure and bleaching in 2020 suggests that directional selection may be already occurring here. Indeed, repeated reductions in coral cover followed by rebounds throughout the last 30 years of disturbance events (**Figure 1**), support the assertion that present-day Keppel coral populations are becoming locally adapted. Evidence of the directional selection toward more heat tolerant S*ymbiodinimum* communities in *A. millepora* populations in this region (Bay et al., 2016; Jones et al., 2008) also supports this hypothesis. Yet, recovery was fastest at sites experiencing the lowest heat stress, and survival was likely enhanced by cooler than average Autumn and Winter temperatures. As climate warming continues, the capacity of corals for recovery is continually being tested (Ortiz et al., 2018; Osborne et al., 2017). Frequent and severe bleaching events, such as the 2020 event reported here, highlight the importance of actions that slow and limit future warming.

## Acknowledgements

We acknowledge the Woppaburra People as the traditional owners of the Keppel Islands in which this research took place. We pay our respects to their elders past, present, and emerging and acknowledge their continuing spiritual connection to their sea Country. All research was done with free prior and informed consent (FPIC) from the Woppaburra Traditional Use of Marine Resources Association (TUMRA) committee were permitted under the Great Barrier Reef Marine Park Authority (GBRMPA) permits G19/43148.1. We thank the crew of the R.V. Cape Ferguson, C. Alessi, K. Allen, D. Brighton, S. Gardener, A. Jones, B. Stephenson, North Keppel Island Environmental Education Centre, and Keppel Dive for assistance in the field. We acknowledge A. Thompson and all present and past members of the AIMS Marine Monitoring Program, as well as the Queensland Parks and Wildlife Service, for their contributions to the historical coral cover data we used here. Thanks to M. Logan for advice on statistical analyses and G. Coleman for assistance with data management. This research was funded by the BHP – AIMS Australian Coral Reef Resilience Initiative.

## Author contributions

CAP, LKB and CJR conceived the study. CAP, CG and CJR collected the data. CAP and CJR analysed the data. CAP, LKB and CJR planned and wrote the paper. CG reviewed and edited the paper. All authors read and approved the final version.

## Footnotes

1 www.ereefs.aims.gov.au

2 www.ereefs.aims.gov.au

