## Supplementary Figure 1 for "High survival following bleaching highlights the resilience of a highly disturbed region of the Great Barrier Reef"

Supplementary Figure 1. Proportion of *Acropora* cover bleached bright white, pale or normal in pigmentation at photo-transect sites in April, June and October 2020. Site abbreviations are defined in Supplementary Table 1**.**

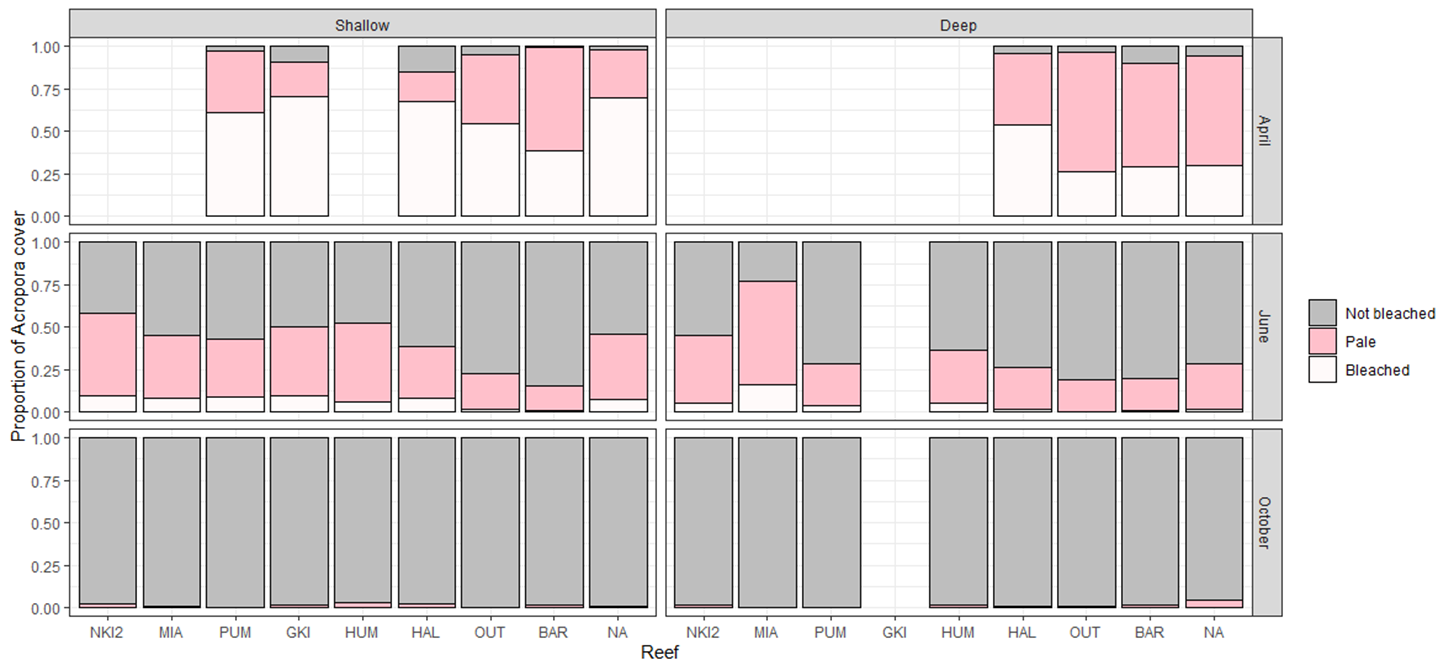
