## Supplementary Figure 2 for "High survival following bleaching highlights the resilience of a highly disturbed region of the Great Barrier Reef"

Supplementary Figure 2. Relationships between maximum daily temperatures and cumulative degree heating weeks (DHW) and median flow rates in east-west and north-south directions. Asterisks indicate statistically significant trends and grey shading represents the 95% confidence intervals around the models.

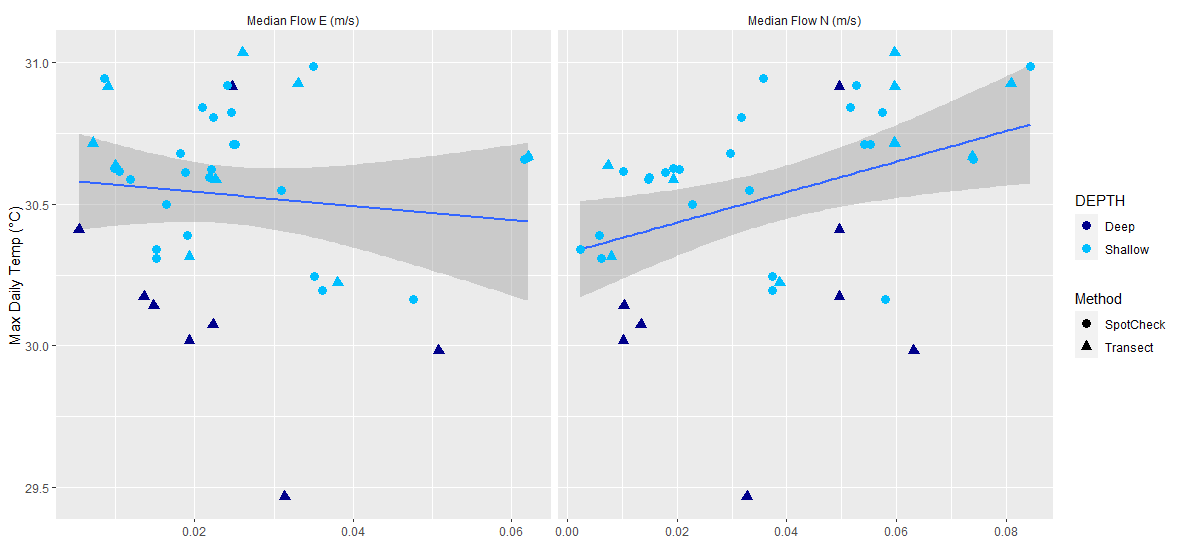

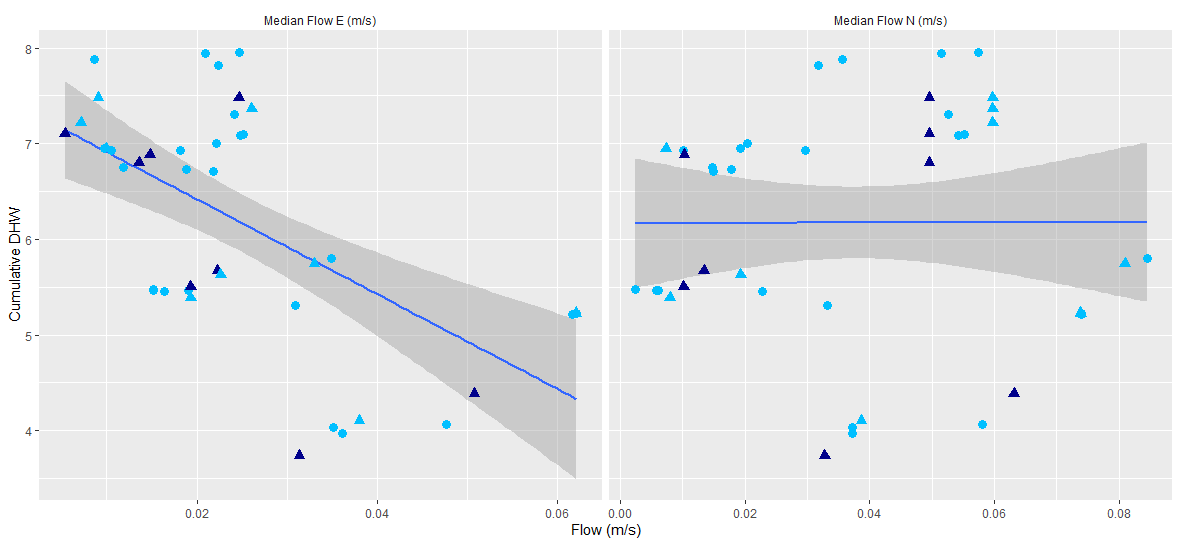

*

*
