## Supplementary Table 1 for "High survival following bleaching highlights the resilience of a highly disturbed region of the Great Barrier Reef"

Supplementary Table 1. Summary of photo-transect sites surveyed in April, June and October 2020 and spot check sites surveyed June 2020 in the Keppel Islands, Great Barrier Reef. *Site locations shown on **Figure 2E**.

| **Date of survey** | | |  |  |  |  |  |
| --- | --- | --- | --- | --- | --- | --- | --- |
| **April 2020** | **June 2020** | **October 2020** | **Depth category** | **Site name**  **(Site abbreviation)** | **Site number*** | **Latitude** | **Longitude** |
| **Photo-transect sites** | | | | | | | |
| 10 April | 24 June | 8 October | Shallow | Pumpkin Island (PUM) | 13 | -23.092 | 150.9034 |
| 10 April | 24 June | 8 October | Deep | Pumpkin Island (PUM) | 13 | -23.092 | 150.9034 |
| 12 April | 22 June | 18 October | Shallow | Outer Rock (OUT) | 20 | -23.6632 | 150.9511 |
| 12 April | 22 June | 18 October | Deep | Outer Rock (OUT | 20 | -23.063 | 150.9508 |
| 12 April | 25 June | 12 October | Shallow | Great Keppel Island, Shelving Bay (GKI) | 23 | -23.1906 | 150.9348 |
| 12 April | 26 June | 9 October | Shallow | Halfway Island (HAL) | 30 | 23.20189 | 150.9719 |
| 12 April | 26 June | 9 October | Deep | Halfway Island (HAL) | 30 | -23.3013 | 150.971 |
| 12 April | 25 June | 9 October | Shallow | Barren Island (BAR) | 34 | -23.1634 | 151.0761 |
| 12 April | 26 June | 9 October | Deep | Barren Island (BAR) | 34 | -23.1637 | 151.0757 |
| 21 April | 23 June | 7 October | Shallow | North Keppel Island, Mazie Bay central (NKI1) | 9 | -23.0855 | 150.9062 |
| 21 April | 23 June | 7 October | Deep | North Keppel Island, Mazie Bay central (NKI1) | 9 | -23.0855 | 150.9062 |
|  | 23 June | 7 October | Shallow | North Keppel Island, Mazie Bay west (NKI2) | 8 | -23.0854 | 150.897 |
|  | 23 June | 7 October | Deep | North Keppel Island, Mazie Bay west (NKI2) | 8 | -23.0854 | 150.897 |
|  | 26 June | 11 October | Shallow | Humpy Island (HUM) | 25 | -23.2151 | 150.9636 |
|  | 26 June | 11 October | Deep | Humpy Island (HUM) | 25 | -23.2151 | 150.9636 |
|  | 26 June | 17 October | Shallow | Miall Island (MIA) | 21 | -23.1551 | 150.9038 |
|  | 26 June | 17 October | Deep | Miall Island (MIA) | 21 | -23.1551 | 150.9038 |
| **Spot-check sites** | | | | | | | |
|  | 22 June |  | Shallow | Conical Island | 1 | -23.0487 | 150.8815 |
|  | 22 June |  | Shallow | Corroboree Island | 2 | -23.0505 | 150.884 |
|  | 22 June |  | Shallow | North Keppel Island, north-west | 3 | -23.0534 | 150.8912 |
|  | 22 June |  | Shallow | Outer Rock, north | 18 | -23.0626 | 150.952 |
|  | 22 June |  | Shallow | Outer Rock, south | 19 | -23.065 | 150.9537 |
|  | 23 June |  | Shallow | North Keppel Island, west | 4 | -23.062 | 150.8887 |
|  | 23 June |  | Shallow | North Keppel Island, south-west | 5 | -23.0801 | 150.8842 |
|  | 24 June |  | Shallow | Square Rock, east | 6 | -23.0993 | 150.8861 |
|  | 24 June |  | Shallow | Square Rock, west | 7 | -23.0996 | 150.8848 |
|  | 24 June |  | Shallow | North Keppel Island, south | 10 | -23.0876 | 150.9114 |
|  | 24 June |  | Shallow | Pumpkin Island, north-east | 11 | -23.0887 | 150.9057 |
|  | 24 June |  | Shallow | Pumpkin Island, north | 12 | -23.0898 | 150.9044 |
|  | 24 June |  | Shallow | Sloping Island, north-east | 14 | -23.0977 | 150.9008 |
|  | 24 June |  | Shallow | Sloping Island, south-east | 15 | -23.1006 | 150.9024 |
|  | 24 June |  | Shallow | Sloping Island, north-west | 16 | -23.0973 | 150.8959 |
|  | 24 June |  | Shallow | Sloping Island, south-west | 17 | -23.1008 | 150.8982 |
|  | 25 June |  | Shallow | Great Keppel Island, North Shelving Bay | 22 | -23.1869 | 150.9337 |
|  | 25 June |  | Shallow | Barren Island, north-east | 31 | -23.1542 | 151.0761 |
|  | 25 June |  | Shallow | Barren Island, north-west | 32 | -23.158 | 151.0697 |
|  | 25 June |  | Shallow | The Child Reef | 33 | -23.155 | 151.0839 |
|  | 27 June |  | Shallow | Bald Rock | 26 | -23.1704 | 150.993 |
|  | 27 June |  | Shallow | Great Keppel Island, near Bald Rock | 27 | -23.1763 | 150.9906 |
|  | 27 June |  | Shallow | Great Keppel Island, Clam Bay west | 24 | -23.1915 | 150.9626 |
|  | 27 June |  | Shallow | Great Keppel Island, Clam Bay central | 28 | -23.186 | 150.9719 |
|  | 27 June |  | Shallow | Great Keppel Island, Clam Bay east | 29 | -23.188 | 150.9764 |
