## Supplementary Table 2 for "High survival following bleaching highlights the resilience of a highly disturbed region of the Great Barrier Reef"

**Supplementary Table 2**. Results of linear models investigating the relationships between four environmental predictors and bleaching prevalence in *Acropora* in April and June 2020 in the Keppel Islands, Great Barrier Reef. Results of Moran’s I test for spatial autocorrelation are presented, and where significant, the applied correlation structure is indicated. Significant *p*-values are indicated in bold.

| **Environmental predictor** | **Sample Period** | **Moran's I p-value** | **Applied correlation structure** | **Slope** | **Standard error** | ***t*-value** | ***p*-value** | **F-statistic** | **n** |
| --- | --- | --- | --- | --- | --- | --- | --- | --- | --- |
| Cumulative DHW | April | 0.861 | N/A | 0.006 | 0.010 | 0.586 | 0.574 | 0.343 | 10 |
|  | June | **<0.001** | Spherical | 0.108 | 0.031 | 3.507 | **0.001** | N/A | 42 |
|  | Change from April to June | 0.0833 | N/A | -0.076 | 0.030 | -2.575 | **0.033** | 6.613 | 10 |
| Maximum daily temperature (°C) | April | 0.644 | N/A | 0.008 | 0.029 | 0.281 | 0.786 | 0.079 | 10 |
|  | June | **<0.001** | Rational quadratic | 0.302 | 0.090 | 3.363 | **0.002** | N/A | 42 |
|  | Change from April to June | 0.914 | N/A | -0.211 | 0.083 | -2.53 | **0.035** | 6.4 | 10 |
| Median flow East-West (m s^-1^) | April | **<0.001** | Spherical | -0.152 | 0.742 | -0.205 | 0.843 | N/A | 10 |
|  | June | **<0.001** | Gaussian | -3.399 | 3.038 | -1.119 | 0.270 | N/A | 42 |
|  | Change from April to June | 0.066 | N/A | 3.682 | 2.349 | 1.567 | 0.156 | 2.457 | 10 |
| Median flow North-South (m s^-1^) | April | **<0.001** | Spherical | 0.004 | 0.464 | 0.009 | 0.993 | N/A | 10 |
|  | June | **<0.001** | Gaussian | 0.632 | 1.837 | 0.344 | 0.733 | N/A | 42 |
|  | Change from April to June | **<0.001** | Exponential | -1.257 | 2.020 | -0.622 | 0.551 | N/A | 10 |
